# Does the mid-domain effect shape interaction networks along environmental gradients?

**DOI:** 10.64898/2026.01.28.702219

**Authors:** Pavel Fibich, Sailee P. Sakhalkar, Robert Tropek, Štěpán Janeček, Yannick Klomberg, Ishmeal N. Kobe, Jan E. J. Mertens, Antigoni Sounapoglou, Tom M. Fayle

**Affiliations:** Institute of Entomology, Biology Centre, Czech Academy of Sciences, Branišovská 31, 37005 české Budějovice, Czechia; Institute of Botany, Czech Academy of Sciences, Zámek 1, 252 43 Průhonice, Czechia; Institute of Microbiology, Czech Academy of Sciences, Vídeňská 1083, 14220 Prague, Czechia; Department of Ecology, Faculty of Science, Charles University, Viničná 7, 12843 Prague, Czechia; School of Biological and Behavioural Sciences, Queen Mary University of London, London, UK

**Keywords:** interspecific interactions, ecological communities, neutral processes, pollination interactions, ant-plant networks, elevational gradients

## Abstract

The mid-domain effect (MDE) predicts that geometric constraints drive unimodal species richness patterns within bounded gradients. However, the role of this effect in ecological networks is currently unexplored. Here we evaluate the role of the MDE in structuring interaction networks. We combine null-model simulations and empirical analyses of plant-pollinator and ant-plant networks along elevational gradients to assess whether the MDE can drive systematic variation in network structure. Our simulations demonstrated that the MDE alone can generate unimodal/U-shaped patterns in network metrics such as connectance, generality, and vulnerability. However, empirical networks only partially conformed to MDE predictions, with deviations indicating the likely influence of other ecological processes. MDE-based models best explained patterns in network-level specialization and nestedness, while only partially explaining patterns in connectance and generality. Because MDEs can shape interaction networks, MDE null models should be used when quantifying the influence of other ecological processes on network structure.

## Introduction

The quantification of ecological networks of interactions between species is critical for understanding ecosystem dynamics. A key focus has been exploring how network structure varies along environmental or geographic gradients, where abiotic factors such as temperature, precipitation, and productivity shape communities (Pellissier *et al*. 2018; Thuiller *et al*. 2024; Tylianakis & Morris 2017). This perspective is especially relevant in the context of ongoing anthropogenic change, which is rapidly altering ecosystems worldwide (Amarasekare 2024). The assumption of many of these studies is that different species have differing environmental requirements, and that this drives variation in network structure. However, the expected null patterns in network structure in the absence of differing environmental niches between species remain unknown. The mid-domain effect (MDE; Colwell & Hurtt 1994), which does not assume differences between species in their environmental niches, has helped to explain patterns in species richness, but has not yet been explored as a driver of patterns in network structure.

The MDE predicts that when species ranges are randomly placed within a bounded domain, such as along an elevational gradient, they will overlap more towards the centre of the domain, generating a unimodal peak in species richness independent of environmental factors (Colwell & Hurtt 1994; Colwell & Lees 2000). Fundamentally, this is because species with longer ranges are constrained to be present in the middle of the domain. If species richness peaks in the mid-domain due to such geometric constraints, corresponding shifts in network structure should be expected, given that species co-occurrence is a prerequisite for interactions and that species richness strongly influences network metrics (Rasmann *et al*. 2014; Vázquez *et al*. 2009). Although this geometric constraint has been extensively studied in species richness patterns (Jetz & Rahbek 2001, 2002; Lees *et al*. 1999), and simulations indicate that food chain length is predicted to be maximal in the mid domain (Prillwitz & Blasius 2020), the broader role of the MDE in structuring ecological networks remains empirically unexplored.

Various bipartite network metrics are expected to exhibit systematic variation along environmental gradients if the MDE operates, because both overlap between species ranges and species richness influence network structure (**Fig. 1**). These predictions are independent of any mechanism involving assumptions about differences between species, and hence have the potential to be used as a minimal null model for explaining variation in network structure along environmental gradients. In particular, the variation in species richness that is generated from MDE models is expected to strongly influence network structure (MacArthur 1968). Previous work has documented relationships between network characteristics and species richness using empirical data and null models, in which species interactions are assigned randomly, with some empirical network metrics following theoretical expectations while others showing deviations. For example, connectance, defined as the proportion of realized interactions relative to all possible links, was observed to decline with increasing species richness, as the number of possible interactions grows faster than the number of realized interactions due to increasing species-specialization (Jordano 1987; Olesen & Jordano 2002; Thébault & Fontaine 2010). Similarly null models predict that connectance will consistently decline with increasing species richness (Blüthgen *et al*. 2008). On the other hand, network generality and vulnerability, measured as the mean number of interaction partners for higher- and lower-level species, respectively, are expected to increase with species richness because the predicted rise in specialization does not outweigh the increasing probability of interactions occurring in species-rich networks, as these two metrics do not correct for network size (Blüthgen *et al*. 2008). In contrast, network-level specialization (*H*_*2*_*’*), is a standardised measure of specialization that accounts for network size (Blüthgen *et al*. 2006), and should consequently not be influenced by species richness, and hence is predicted not to show MDE driven patterns along environmental gradients. Nestedness, which quantifies the tendency of specialists to interact with subsets of generalist partners, tend to increase with species richness in the empirical networks (Bascompte *et al*. 2003; Vázquez & Aizen 2004; Vázquez & Stevens 2004). However, nestedness in null models does not consistently increase with species richness, but rather peaks at intermediate species richness (Bascompte *et al*. 2003; Rodríguez-Gironés & Santamaría 2006).

**Fig. 1.**
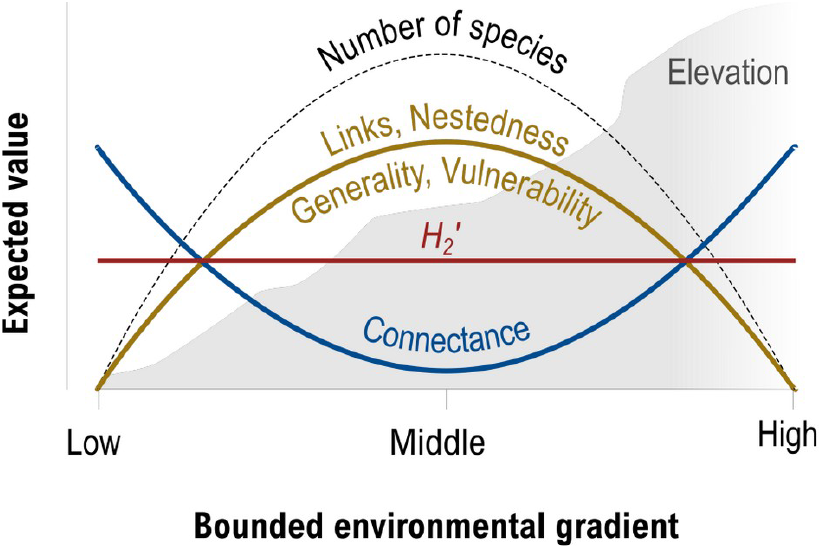
Expected trends for species richness and network metrics along a bounded gradient (e.g. elevation), following mid-domain constraints. The total realized links, nestedness, generality, and vulnerability (yellow solid line) are expected to show a unimodal trend following the mid-domain effect in species richness (black dashed line). In contrast, connectance is expected to show a U-shaped pattern (blue solid line), whereas no systematic trend is expected for *H2′* (red solid line). See main text for motivations underlying these hypothesised trends.

Because network metrics are expected to vary along environmental gradients in this way, even in the absence of differences between species in environmental niches or their biotic interactions, any theory that makes predictions based on such differences should be evaluated against a null MDE network model. For example, MacArthur (1968) and Dobzhansky (1950) hypothesised that species in more species-rich communities tend to exhibit narrower niches due to increased competition and long-term stability, respectively, both leading to higher specialization. These predictions were formalised into the latitude-niche breadth hypothesis (MacArthur 1968) and later extended to elevational gradients (Rasmann *et al*. 2014). Observed patterns in network structure along environmental gradients are likely to result from a combination of network MDEs and niche-based drivers such as these. Indeed, empirical studies reveal complex and sometimes contrasting patterns in network structure and species richness along environmental gradients. For example, as predicted by null modelling, connectance, generality, and vulnerability declined with increasing species richness in a large-scale study using 65 food webs of various types (Riede *et al*. 2010) and in a separate analysis of 54 pollination networks (Trøjelsgaard & Olesen 2013). However, Schleuning *et al*. (2012) found the opposite pattern, with these three network metrics increasing with species richness in an analysis of 282 plant-animal mutualistic networks. Similarly, while nestedness increased with species richness in plant-pollinator networks (Nielsen & Bascompte 2007; Trøjelsgaard & Olesen 2013), no such relationship was found in ant-plant networks (Dáttilo & Vasconcelos 2019). In contrast, the relationship between network-level specialization (*H*_*2*_*’*) and species richness is highly inconsistent. While some studies found a positive relationship (Dalsgaard *et al*. 2011), others negative correlations (Classen *et al*. 2020; Schleuning *et al*. 2012), some found no significant relationship (Dáttilo & Vasconcelos 2019; Sakhalkar *et al*. 2025). These discrepancies indicate that while species richness plays a fundamental role in structuring interaction networks, with a potential contribution of MDEs, additional ecological drivers are likely to be important. While numerous studies have explored how various environmental and ecological variables shape ecological networks, the role of the MDE remains largely overlooked, highlighting the need for further investigation to better understand the complex drivers of interspecific interactions. Without quantification of the null expectations under MDE assumptions, evaluation of any niche-driven hypotheses relating network structure to the environment is likely to be highly challenging.

Here, we analyse whether mid-domain effects (MDEs) drive variation in bipartite network structure along environmental gradients by combining simulation-based null models and empirical network analyses. Our study represents the first evaluation of MDEs in the context of ecological networks, providing a direct link between geometric constraints and network patterns. Based on theoretical predictions (**Fig. 1**), we expect the unimodal pattern of species richness related to MDE to generate unimodal or U-shaped patterns of network metrics along environmental gradients. Using simulation models, we test whether the random placement of species ranges within a bounded domain can generate such patterns in connectance, network-level specialization (*H*_*2*_*’*), generality, vulnerability, and nestedness. We further investigate how MDE-driven patterns are influenced by species’ range sizes and the number of interacting partners. Additionally, we compare empirical bipartite interaction networks sampled along elevational gradients to MDE null models in which observed species ranges are randomly shifted along the gradient, allowing us to determine the degree to which mid-domain effects explain observed patterns in network structure.

## Methods

To test whether mid-domain effects (MDE) drive variation in bipartite network structure along environmental gradients, we combined simulations with empirical data analysis. First, we developed a null mid-domain model that simulates bipartite interaction networks based on random species range placement within a bounded domain, incorporating different levels of species specialization and range sizes (**Fig. 2**). This allowed us to assess whether MDE alone can generate unimodal or U-shaped patterns in network metrics such as connectance, and nestedness (**Fig. 1**). Second, we applied this null model to four empirical datasets of plant-pollinator and ant-plant networks recorded along elevational gradients to test whether observed network patterns align with MDE expectations, identifying cases where mid-domain constraints were not sufficient to explain observed network patterns, hence indicating likely presence of other drivers of network structure.

**Fig. 2.**
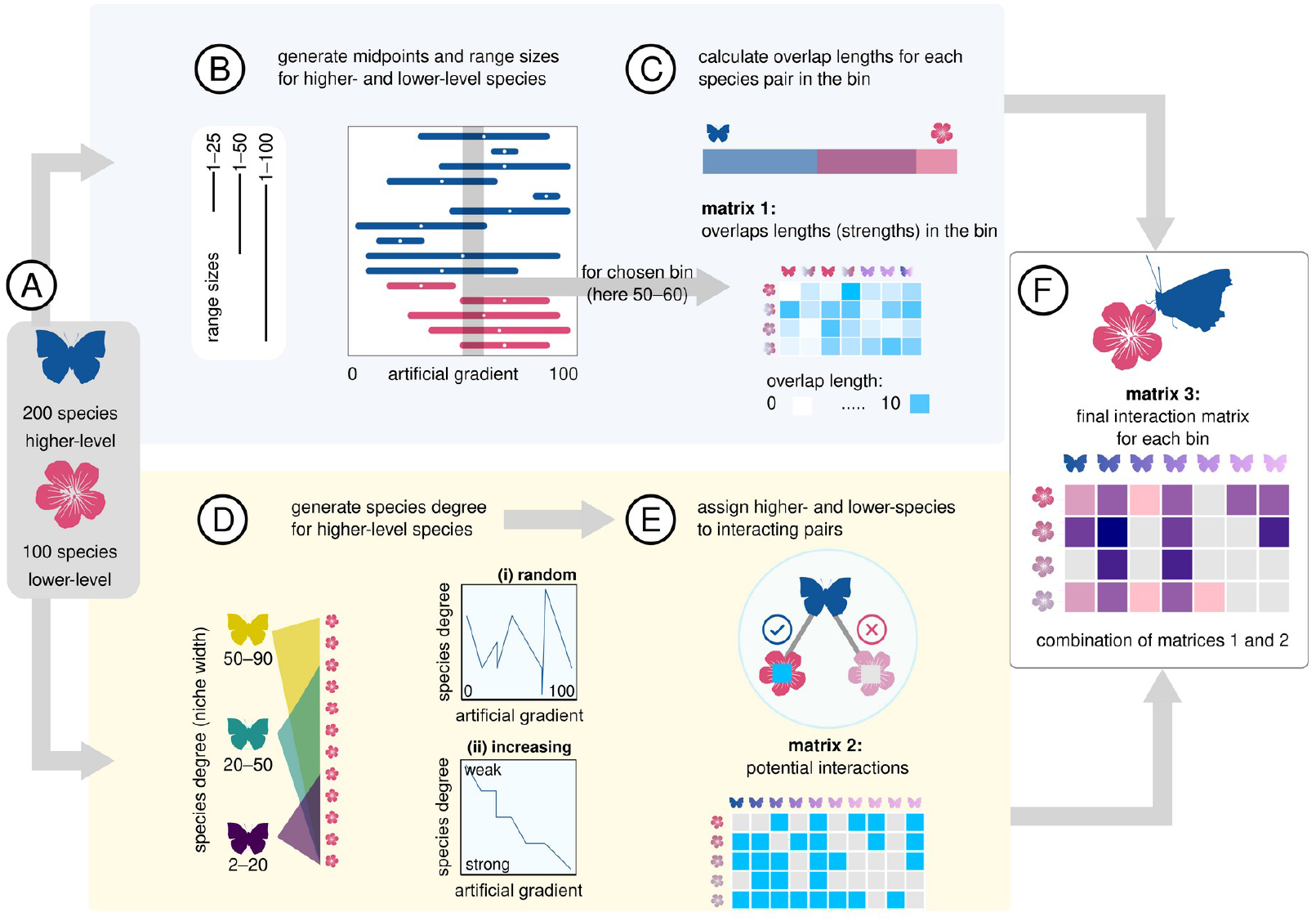
Overview of null model construction. Butterfly icons represent higher-level species, and flower icons represent lower-level species. (A) The model was initialized with 200 higher-level species and 100 lower-level species. Blue background: (B) Species ranges and midpoints were generated along a one-dimensional artificial gradient of 100 arbitrary units, divided into bins of 10. The 50–60 bin is highlighted in grey as an example of a bin with overlapping species” ranges. The model simulated three species range scenarios: 1–25, 1–50, and 1– 100 units. (C) Species range overlaps for the selected bin were calculated and recorded in an **overlap strength matrix** (**matrix 1**). Yellow background: (D) Higher-level species were assigned a species degree (number of lower-level interaction partners) within three categories: generalists (50–90 partners, purple), intermediate specialists (20–50 partners, green), and specialists (2–20 partners, yellow). Species degree was simulated under two scenarios: random species degree, and systematically increasing species degree along their mid points towards one end of the gradient, representing higher specialization. (E) Lower-level species were randomly assigned as potential partners based on the species degree constraints of higher-level species, creating a **potential interaction matrix** (**matrix 2**). (F) The three matrices (overlap strength, species co-occurrence, and potential interactions) were combined to produce a **final quantitative interaction matrix** (**matrix 3**) for each bin, which was used to calculate network indices. This process was repeated 100 times per scenario, ensuring robust estimates of network properties across different simulation scenarios.

### Interaction network simulations

To explore whether MDE can generate predictable patterns in network structure, we simulated the assembly of species interaction networks along a one-dimensional artificial gradient (see **Fig. 2**. and **Fig. S1** for an overview, and **Table S1** for simulation parameters). This artificial gradient represents a bounded domain (a constrained spatial or ecological range) divided into arbitrary units and does not correspond to a specific environmental variable. The total gradient length was set to 100 arbitrary units. Within this domain, species were assigned continuous ranges spanning a subset of the gradient, with midpoints randomly placed along its length. By modelling species distributions and interactions, such as variation in the range length and species degree (number of interaction partners), in this simplified null modelling framework, we aimed to assess whether geometric constraints alone could shape key network properties.

We began with an artificial species pool, consisting of 100 lower-level species (representing e.g. plants) and 200 higher-level species (representing e.g. pollinators) (**Fig. 2A**). Each species was assigned a continuous range spanning a subset of the gradient, with midpoints randomly placed along the full domain length (1–100 arbitrary units; **Fig. 2B**). To simulate different levels of range restrictions, we modelled three scenarios where range lengths followed uniform distributions of 1–100 (where the widest range could span the entire gradient), 1–50, and 1–25 arbitrary units. If a species’ range extended beyond the domain limits, we generated a new midpoint at a random location on the gradient to ensure it remained entirely within the gradient, following Model 2 in Colwell and Hurtt (1994). All modelled scenarios determined species overlap along the gradient, which is a prerequisite for species to interact. All scenarios resulted in mid-domain species-richness peaks due to greater species overlap at the centre of the environmental gradient, with shorter species ranges producing less pronounced peaks (examples in **Fig. S2**), consistent with previous findings (Colwell *et al*. 2009).

Once the range sizes and placements were simulated, we divided the gradient into ten equal bins (segments of 10 arbitrary units: 1–10, 11–20, …, 91–100). Using other bin lengths did not affect the results substantially. For each bin, we calculated the extent of the range overlap for each species pair between 0 and 10, which was recorded in an **overlap strength matrix** (**Matrix 1; Fig. 2C**). To define interactions between higher-level and lower-level species (e.g., if a given pollinator species visits flowers of a plant species), we assigned species degrees to higher-level species, simulating different levels of specialization in their relationships. Using a uniform distribution, we generated species-specific values for the number of lower-level species each higher-level species interacted with **(Fig. 1D)**. These values were categorised into three levels, from high to low specialization: (i) specialists, interacting with 2–20 lower-level species, (ii) intermediate specialists, interacting with 20–50 lower-level species, and (iii) generalists, interacting with 50–90 lower-level species. We simulated communities of higher-level species following two scenarios on how specialization varied along the gradient: (i) random specialization, where species degree of higher-level species varied independently of position along the gradient, and (ii) increasing specialization, where randomly generated species degree of higher-level species was sorted to increase along the gradient for increasing species mid-domains, simulating a pattern from weak to strong specialization along the gradient. Note that simulating the reverse pattern, with stronger specialization at lower gradient values, would simply produce a mirrored response, given that the gradient is not directional. Using these species-specific interaction constraints, we randomly assigned lower-level species to interact with higher-level species, generating a **potential interaction matrix** (**matrix 2; Fig. 1E**), which determined which species pairs could interact within each bin along the gradient. This component of our simulation did not include any sort of a priori mid-gradient peak in specialization. The **final quantitative interaction matrix** for each gradient bin was constructed by combining the three matrices described above (**Fig. 1F**). If a species pair was present in a bin and their interaction was allowed, their value in the interaction matrix (interaction strength) was set to the length of their overlap in the bin. Otherwise, their interaction strength was set to zero (see **Fig. S1** for an example).

For each simulated final quantitative interaction matrix, we computed six common bipartite network metrics representing different aspects of network structure (reviewed in Dormann *et al*. 2009): total realized links (number of non-zero values in the matrix), connectance (proportion of realized interactions relative to all possible links), unweighted nestedness (in particular NODF, a nestedness measure quantifying the extent to which species with fewer interactions form subsets of those with more interactions), network-level specialization *H*_*2*_*’* (a metric that is standardized for network size and species interaction frequencies, ranging from 0 for highly generalised networks to 1 for highly specialized ones; Blüthgen et al. 2006), generality (weighted mean number of lower-level species each higher-level species interacts with), and vulnerability (weighted mean number of higher-level species each lower-level species interacts with). All network metrics were computed using the ‘bipartite’ R package (Dormann *et al*. 2022).

We simulated all scenario combinations using 500 replications each and reported the mean values for each network metric. To test whether the simulated network metrics exhibited unimodal/U-shaped, more complex or linear patterns along the gradient, we compared second- and fourth-degree polynomial fits to linear fits using F-statistics (P < 0.05 and delta AIC>2). We constructed the models using the *lm* function in the “stats” R package and compared model fits for each metric using ANOVA tests with the *anova* function. The full simulation code is available at https://github.com/pavel-fibich/mdefood.

### Empirical interaction network datasets

To test whether the MDE could in part explain the structure of real-world bipartite interaction networks, we analysed four empirical plant-animal interaction networks recorded along elevational gradients **(Table 1)**. Elevational gradients provide an ideal system for testing MDE because they are bounded at both ends and exhibit predictable environmental variation, including temperature, precipitation, and vegetation structure, over short spatial scales (Colwell & Hurtt 1994). Importantly, elevation allows quantification of the mid-domain effect on a single dimension, simplifying the analysis and interpretation of species richness and network structure patterns (Colwell & Hurtt 1994). This bounded one-dimensional gradient allows for clearer comparisons between observed and expected patterns under mid-domain models. We used four empirical datasets (for more detail see Supplementary Text S1): a tropical myrmecophytic ant-plant network from Papua New Guinea (Plowman *et al*. 2017), a tropical dry-season plant-pollinator network from Cameroon (unpublished), a tropical wet-season plant-pollinator network from Cameroon (unpublished), and a temperate plant-pollinator network from Czechia (unpublished).

**Table 1.** Details of the four empirical interaction networks sampled along elevational gradients. *The sparse networks from the three highest elevations for this dataset were merged, as were the three other pairs of adjacent elevations, resulting in four merged networks.

| Empirical dataset and reference | Location | Gradient range (m a.s.l.) | Number of sampled elevations | Higher-level species richness, taxon | Lower-level species richness, taxon |
| --- | --- | --- | --- | --- | --- |
| **Myrmecophytic ant-plant dataset** (Plowman *et al.* 2017) | **Mount Wilhelm, Papua New Guinea** | 700–1400 | 9, reassembled to 4 bins* | 10, ants | 21, trees |
| **Temperate plant-pollinator dataset** (unpublished) | **Krkonoše Mountains, Czechia** | 450–1000 | 4 | 45, insects | 26, herbs |
| **Tropical wet-season plant pollinator dataset** (unpublished) | **Mount Cameroon, Cameroon** | 650–2200 | 4 | 161, animals | 81, plants |
| **Tropical plant dry-season pollinator dataset** (unpublished) | **Mount Cameroon, Cameroon** | 650–2200 | 4 | 401, animals | 121, plants |

To standardise resolution across the datasets, we combined sparse data from adjacent sampled elevations in the ant-plant dataset, ensuring that all four datasets had four elevational bins (domains, **Table 1**). For each dataset, we determined the range of all lower- and higher-level species based on their observed occurrences in recorded networks along the elevational gradient. For species with discontinuous distributions in sampled networks, we assumed continuous ranges.

To generate null MDE models, we randomly shifted species’ ranges along the elevational gradients (n = 500 simulations per dataset), ensuring that the entire range of each species remained within the bounded domain. For example, in a four-bin elevational gradient, a species recorded at bins 1 and 2 could either remain at these bins or be shifted to bins 2 and 3 or 3 and 4, with each option being equiprobable. This shuffling algorithm is equivalent to that used in the mid-domain null model simulations (see above). Species were considered able to interact if their ranges overlapped in a given bin. Interaction strength was assigned based on observed interaction abundance across all elevations. For the ant-plant dataset, interaction strength was the frequency of ant inhabitation of each ant-plant (myrmecophyte) species, while for the plant-pollinator datasets, it was the frequency of visits per flower of each plant species. For every observed and randomised network, we calculated the same network structure metrics as in the simulated datasets (see above). These calculations were performed separately for each of the four elevation bins. To assess whether observed networks deviated from null expectations, we calculated a measure based on the sum of standardised effect sizes (SES = (observed - mean null)/SD null) for each metric across all elevations for that dataset. Observed interaction network characteristics were considered to deviate significantly from the MDE model for a particular dataset if the sum of the observed SES values exceeded 95% of the sum of the SES values from randomised networks.

## Results

### Interaction network simulations

Most network metrics exhibited expected unimodal or U-shaped relationships along the artificial gradient (**Figs. 3, 4**), consistent with the mid-domain effects expectations (**Fig. 1**). Higher- and lower-level species richness followed a unimodal pattern regardless of species range span, specialization degree (from low to high) or whether specialization changed directionally along the gradient. Total realized links, generality, and vulnerability showed mostly unimodal patterns, consistently peaking at the gradient mid-point. Network-level specialization (*H*_*2*_*’*), nestedness and connectance showed mostly U-shaped patterns with the lowest values at the gradient mid-point and more flat patterns for nestedness and connectance than for *H*_*2*_*’*, except under increasing specialization scenarios.

**Fig. 3.**
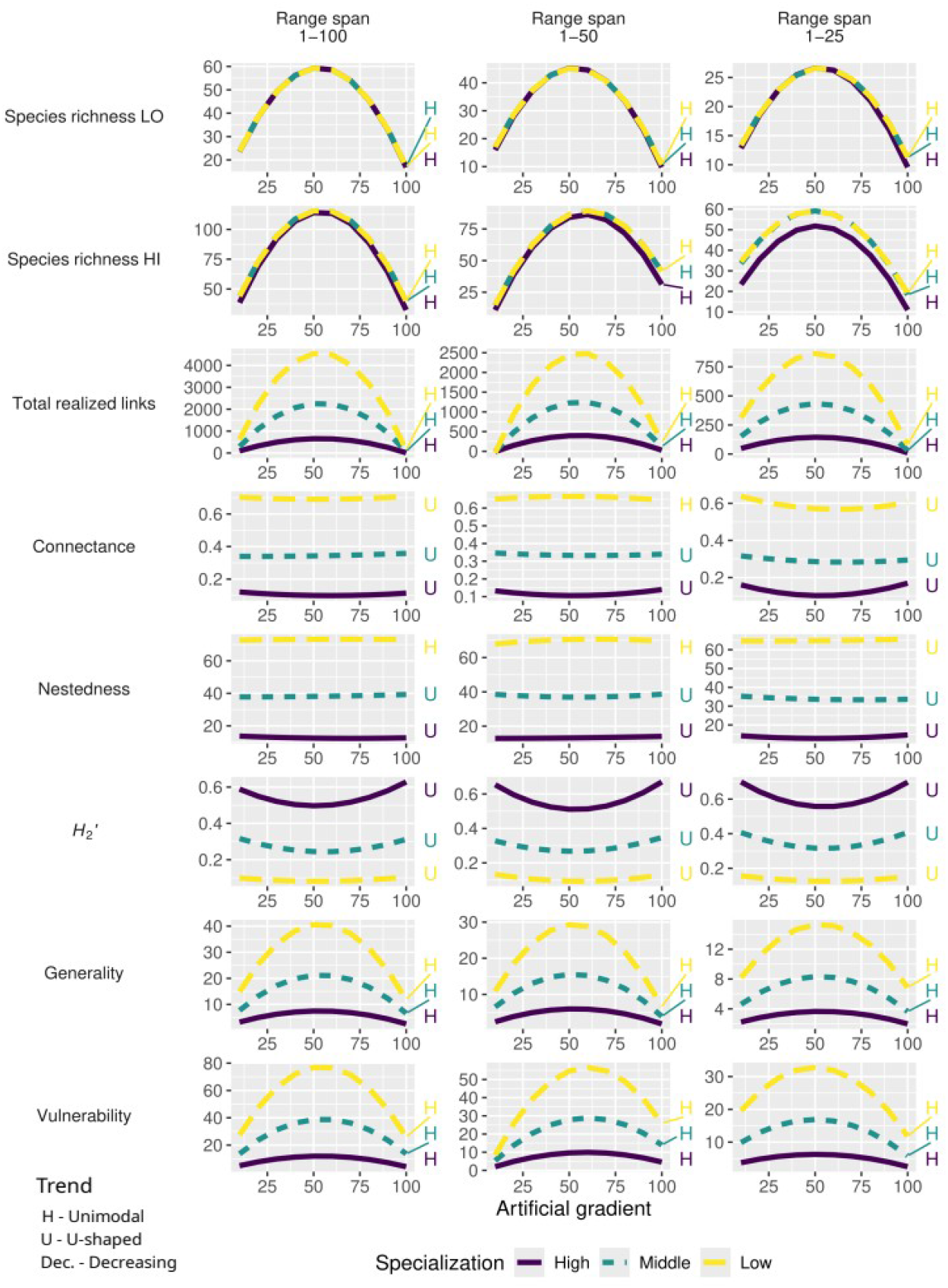
Predicted changes in network structure under mid-domain null models where higher-level species’ specialization degree is randomly distributed along the artificial gradient. Panel columns represent modelled scenarios with different species’ range sizes, from wide-ranged (left) to short-ranged (right) species. Panel rows correspond to network metrics. Within each panel, the best-fitting models are shown for simulations with the three categories of modelled species degree: yellow dashed lines represent low specialization with generalists in the community (species degree 50–90), green short-dashed lines represent middle specialization with intermediate specialists (20–50), and purple solid lines represent high specialization with specialists (2–20). The second-degree polynomial trends of individual lines are described by letters: H – unimodal (hump-) shaped, and U – U-shaped.

**Fig. 4.**
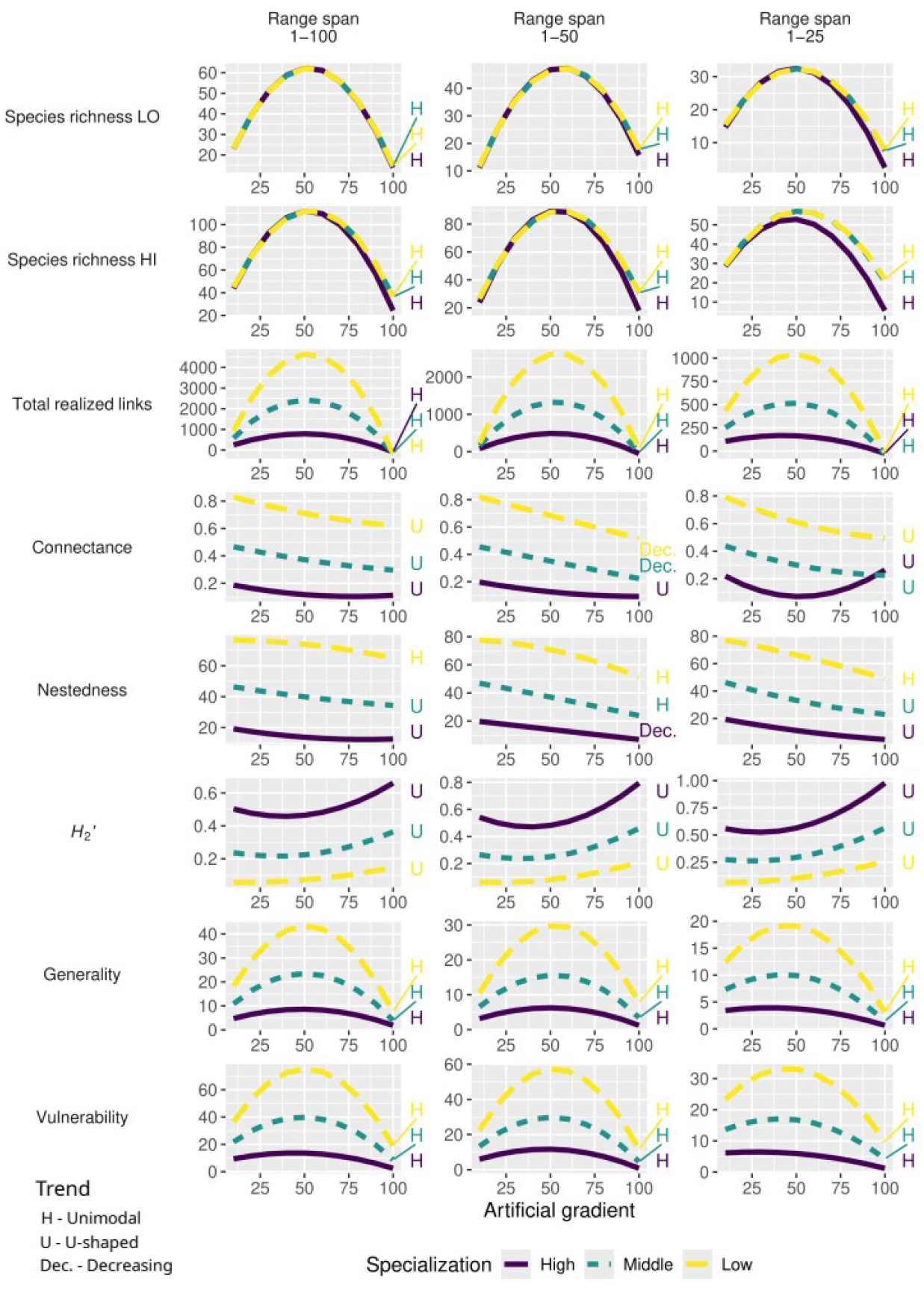
Predicted changes in network structure under mid-domain models where higher-level species” specialization degree systematically increases along the artificial gradient. Panel columns represent modelled scenarios with different species” range sizes, from wide-ranged (left) to short-ranged (right) species. Panel rows correspond to network metrics. Within each panel, the best-fitting models are shown for simulations with the three categories of modelled species degree: yellow dashed lines represent low specialization with generalists in the community (species degree 50–90), green short-dashed lines represent middle specialization with intermediate specialists (20–50), and purple solid lines represent high specialization with specialists (2–20). The second-degree polynomial trends of individual lines are described by letters: H – unimodal (hump-) shaped, U – U-shaped, and Dec. – linearly decreasing.

When specialization was distributed randomly along the gradient (**Fig. 3**), *H*_*2*_*’* displayed symmetric U-shaped patterns across all range-size scenarios. Connectance and nestedness mostly followed similar U-shaped patterns, expect for networks with low specialization, and medium-sized range spans, where connectance and nestedness both showed a unimodal pattern. Generality and vulnerability showed unimodal distributions, mirroring the pattern in the number of interactions.

When simulated specialization increased systematically along the gradient (**Fig. 4**), distinct shifts in network metrics emerged. *H*_*2*_*’* showed a left-skewed U-shaped pattern. Connectance exhibited mostly a right-skewed U-shaped pattern, contrasting with the symmetric U-shaped patterns observed under models with random specialization distributions. Generality and vulnerability maintained unimodal distributions. Nestedness predominantly followed right-skewed U-shaped patterns, except for low specialization, which exhibited right-skewed unimodal or decreasing trends.

The more complex, the fourth-degree polynomial curve fitting mostly showed similar patterns to the second-degree polynomials (**Figs. S3 and S4**).

### Empirical interaction networks

Most observed network metrics fell within the 95% confidence interval of the distribution predicted using randomised species’ distributions along the elevational gradients (**Figs. 5** and **S6**). Of the 128 individual combinations of elevations and network metrics for individual empirical datasets, 79 values (62%) fell within the 95% confidence interval of the null model predictions. Nonetheless, all empirical interaction networks showed some deviations from the null model predicted values for at least one of the network metrics.

**Fig. 5.**
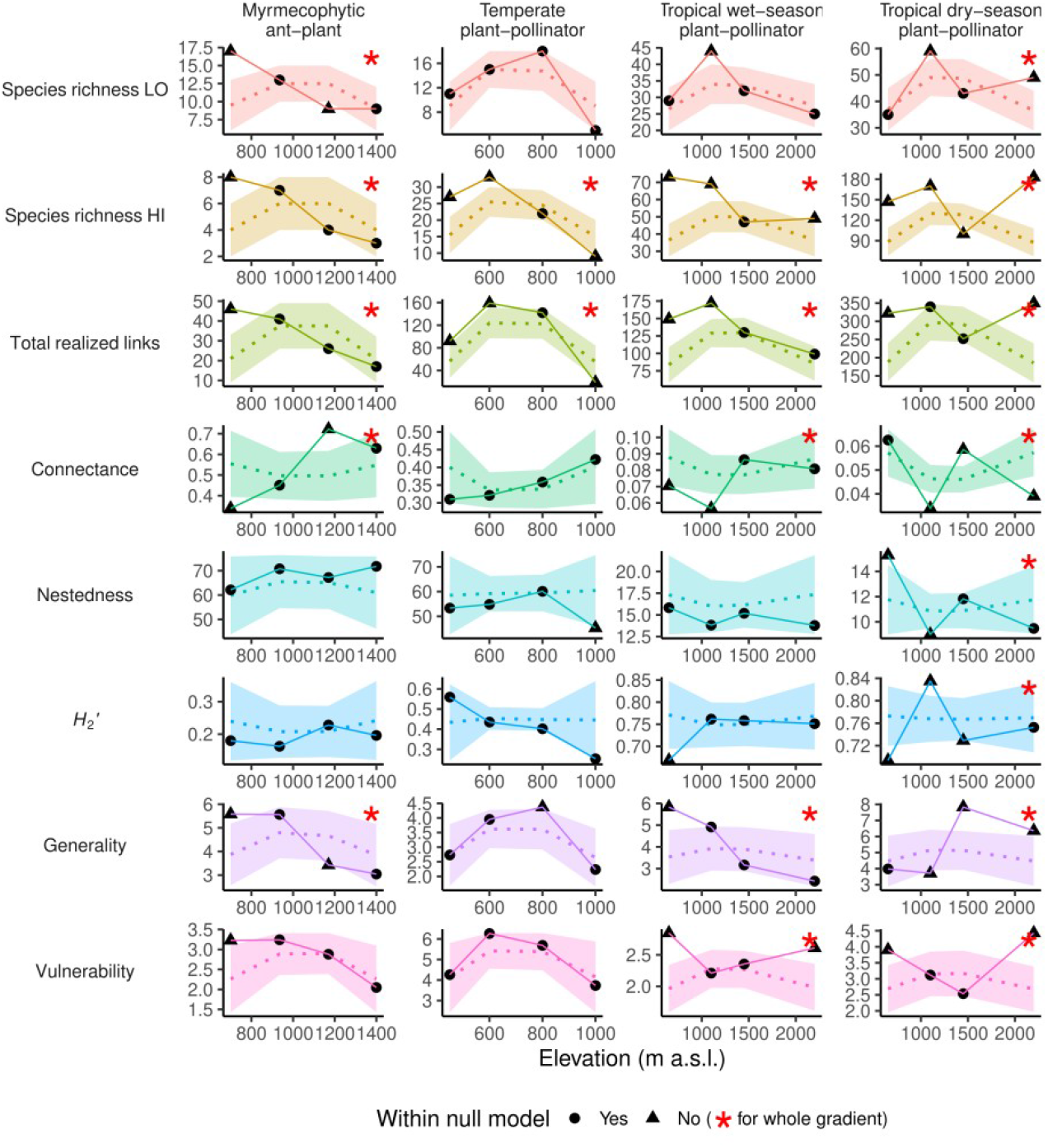
Patterns of network metrics in empirical interaction networks along elevational gradients, compared to predictions from mid-domain network null models (networks with randomised species” distribution along the elevational gradients). Solid lines represent observed patterns, dotted lines denote null model means, and coloured areas delimit the 95% confidence intervals of null model values. Empty circles and solid triangles indicate whether the observed network metric for a given elevation fall within or outside of 95% confidence intervals of null model values, respectively. Asterisks (*) in the top-right corner of each panel indicate network metrics where the sum of observed SES values along the whole gradient exceeds 95% of the sums of SES values from null models, signifying that the overall pattern along the gradient significantly deviates from random expectations for that network metric.

When considering model fits for entire elevational gradients, 12 out of 32 patterns of empirical network metrics followed patterns predicted by the MDE null models, whereas the remaining metric/network combinations deviated to varying degrees. Nestedness and *H*_*2*_*’* were the best-fitting metrics with observed patterns closely matching MDE null model predictions, whereas patterns of total realized links and higher level species number deviated the most from null-model expectations. The tropical dry-season plant-pollinator networks exhibited the highest deviations from the null models, with all network metric patterns falling outside the 95% confidence intervals. In contrast, the temperate plant-pollinator dataset most closely followed the null models, with only total realized links and higher level species number deviating slightly from the null model predictions.

## Discussion

Our work provides important insights into how mid-domain effects (MDE) influence interaction network structure. Our simulations under MDE assumptions alone generated predictable unimodal/U-shaped patterns in key network metrics, such as total realized links, connectance, and generality, and vulnerability. However, our empirical results revealed some deviations from MDE expectations, particularly in total realized links, connectance and generality, suggesting that additional factors play a role in structuring real-world networks.

### Mid-domain effects in network structure: Evidence from null models

Our null modelling approach confirmed that MDE alone can generate predictable patterns in network structure. Randomly distributed species ranges along the gradient resulted in the mid-domain peaks in species richness (**Fig. 1**), recapitulating the results of previous studies (Colwell & Hurtt 1994; Colwell & Lees 2000).

Total realized links, generality, and vulnerability followed unimodal patterns, with the highest values occurring at mid-gradient, where species richness also peaked, while connectance followed a U-shaped pattern, with the lowest value in the middle of the gradient. These results align with theoretical expectations from previous null models that demonstrate increasing species richness to directly affect the total realized links and connectance (Blüthgen *et al*. 2008; Olesen & Jordano 2002), and the connectance-dependent metrics generality and vulnerability (Blüthgen *et al*. 2008). These MDE effects can be attributed to the strong influence of species richness on species co-occurrence patterns, which can in turn determine interaction opportunities (Vázquez *et al*. 2009).

However, other network metrics did not follow our predictions for their patterns along the artificial gradient, even though species richness followed a unimodal pattern. Nestedness varied with specialization degree and ordering, mostly showing U-shaped patterns, although previous null modelling approaches predict that nestedness should peak at intermediate species richness or show increasing pattern (Bascompte *et al*. 2003; Rodríguez-Gironés & Santamaría 2006; Vázquez & Aizen 2004). However, the magnitude of the changes in nestedness along the gradient were extremely small, relative to the impacts of specialization. Network-level specialization (*H*_*2*_*’*) deviated from the predictions, showing a unimodal pattern that contrasted with the null expectation that specialization should remain independent of network size in null modelled systems without additional ecological constraints (Blüthgen *et al*. 2006). This could be because our simulations used specific rules to determine whether species overlap in their ranges, and if they overlap, whether they interact, and hence specialization can be influenced beyond just the impacts of variation in species richness.

The length of species ranges along the gradient directly affected species richness and hence network size (the total number of species in a network and the total realized links). Although generality and vulnerability reflected these patterns in network size, this is likely because neither metric includes a correction for network size (Blüthgen *et al*. 2006). Overall patterns of network metrics did not depend on the lengths of species ranges in our simulations. In real networks, the length of species ranges is interconnected with species specialization, with broader-ranging species tending to be more generalised than those with restricted ranges (Trøjelsgaard & Olesen 2013). While specialization (if there are more generalists or specialists in the community) was related in our simulations to network size and thus also the absolute values of network metrics, we did not observe any strong effects of specialization on the overall network patterns along the gradient. Neither the length of species ranges nor specialization degree were directionally related to the gradient, and so these parameters tended to affected network size uniformly along the gradient. On the other hand, when we ordered specialization along the gradient (**Fig. 4**), compared to the random distribution of specialists and generalists (**Fig. 3**), this had impacts on connectance, nestedness, and H_2_’, with the latter showing the greatest magnitude of change. As expected, by increasing species level specialization towards the upper end of the artificial gradient, in way that mimicked some environmental gradient, we observed increased network-level specialization, and decreased connectance and nestedness, that is concordant with empirical studies along elevational gradients (e.g. Ramos-Jiliberto *et al*. 2010; Schleuning *et al*. 2012; Hoiss *et al*. 2015). Nonetheless, all three network metrics retained weak U-shaped patterns driven by MDE.

### Empirical networks: mixed support for MDE

Our empirical networks provided mixed support for MDE-driven patterns, with some datasets and network metrics aligning well with MDE predictions, while others showing strong deviations (12/32 empirical metric/dataset combinations showing good fit). However, even those datasets with good fits to MDE predictions did not always show unimodal/U-shaped patterns along the gradient. Nestedness and *H*_*2*_*’* were the most consistent metrics, matching MDE expectations reasonably well, whereas total realized links and higher level species number showed the strongest deviations from MDE models. Other network metrics varied in their fit to MDE models across the empirical datasets. Differences among datasets may potentially be linked to variations in environmental conditions, gradient length, species turnover, and the type of interspecific interactions. These analysed datasets originate from different geographic areas, including the European temperate region, the Afrotropics, and a tropical Pacific island, with distinct evolutionary histories and species pools. The diverse environmental conditions probably resulted in different patterns in interaction specialization, as predicted by the latitude-niche breadth hypothesis (Baiser *et al*. 2019; MacArthur 1972).

Similarly, previous studies reveal more complex and sometimes contrasting patterns of network metrics along elevational gradients. For example, connectance follows U-shaped patterns with elevation in some networks where species richness peaks at mid-elevations, as observed in host-parasite networks (Morris *et al*. 2015) and plant-pollinator (Adedoja *et al*. 2018; Hoiss *et al*. 2015), supporting our theoretical predictions. However, nestedness shows inconsistent patterns along these gradients, varying across systems (Adedoja *et al*. 2018; Minachilis *et al*. 2020). In cases where species richness varies unpredictably along elevational gradients, such as flower-visitor networks in Colombia (Cuartas-Hernández & Medel 2015), connectance still declines with increasing species richness, while nestedness again shows no systematic relationship. Generality and vulnerability decline with elevation in host-parasitoid networks (Maunsell *et al*. 2015), with temperature in plant-herbivore network (Pitteloud *et al*. 2021), and with species-richness in plant-frugivore network (Quitián *et al*. 2018), despite opposing predictions from both ecological theory and null modelling expectations (Blüthgen *et al*. 2008). Network-level specialization shows varying relationships with species richness along elevation, with some studies reporting a decreasing trend (Adedoja *et al*. 2018; Dzekashu *et al*. 2023; Lara-Romero *et al*. 2019), others an increasing trend (Aguirre & Junker 2024; Lei *et al*. 2023), and at least one finding no systematic pattern (Classen *et al*. 2020). The strongly varying patterns across different elevational gradients in our analyses of empirical datasets as well as that found in previous studies, with some network metrics aligning with MDE predictions while others deviating, indicate that although ecological mechanisms are a dominant force shaping interaction networks, their interplay with mid domain effects may be context-dependent.

### Future research directions

While our MDE simulations provide valuable insights into the role of geometric constraints in shaping ecological networks, these models rely on simplifying assumptions that may limit their applicability to real-world systems. Our model assumes that species ranges are uniformly distributed along the gradient, disregarding spatial heterogeneity, species-specific habitat preferences, and asymmetric dispersal, all of which can strongly influence species distributions and interactions (Bahn *et al*. 2006; He *et al*. 2022). One way of accounting for deviations from these null model assumptions it to use mid-point attractors (Colwell *et al*. 2016), where environmental favourability varies along the gradient, potentially skewing mid-domain patterns. Such asymmetrical mid-domain peaks in species richness have been documented for various insects (Dolson & Kharouba 2024; Maicher *et al*. 2020; Toko *et al*. 2023). Additionally, in our simulations, species interactions were assigned based solely on range overlap, without accounting for trait-based partner preferences or interaction strengths (Bartomeus *et al*. 2016), which may contribute to deviations from empirical patterns. Future implementations of the model could incorporate these factors to improve model realism.

Our simulations also explored community configurations consisting of either generalists, intermediate specialists, or specialists. While such configurations are unlikely to occur in nature, they provide useful benchmarks, as some networks are strongly specialist-dominated, such as hummingbird and sunbird pollination networks (Janecek *et al*. 2021; Maglianesi *et al*. 2024), while others are composed primarily of generalists, such as hoverfly pollination networks (Branquart & Hemptinne 2000). Our intermediate scenarios may align with systems of limited specialization, such as plant-frugivore (Schleuning *et al*. 2011) and ant-plant networks (Blüthgen *et al*. 2007). Furthermore, our models assume that species degree distributions are either random or increase systematically along the gradient, whereas empirical networks may exhibit more complex patterns shaped by local ecological interactions, competition, and environmental gradients (reviewed in Vázquez *et al*. 2009). Such systematic variation in specialization could be further explored in the future using our modelling framework.

Finally, while our empirical networks were collected along elevational gradients, our framework can be extended to other environmental and spatial gradients, such as latitude (Colwell & Lees 2000; Lyons & Willig 2002). Although latitudinal species richness gradients are variable but well-documented (reviewed in Willig *et al*. 2003), their effects on network structure remain understudied (Forister *et al*. 2015; Moles & Ollerton 2016; Schleuning *et al*. 2011; Zvereva *et al*. 2024). To our knowledge, the role of MDE in structuring ecological networks along latitudinal or other gradients has never been analysed and should be a focus for future research crucial in understanding the processes structuring communities along large-scale gradients.

## Conclusion

Our study demonstrates that mid-domain effects alone can generate the unimodal or u-shaped patterns in network structure observed empirically along some environmental gradients. Species range size and degree of specialization were found to strongly affect the null MDE predictions. While our null models predicted unimodal/U-shaped patterns in key network metrics, real-world networks showed strong deviations from these expectations, particularly in species-rich systems. This indicates that other ecological processes, such as environmental filtering and kinds of interspecific interactions other than those modelled, also play a dominant role in structuring interaction networks. This highlights the importance of integrating MDEs and niche-based processes when interpreting network structure along environmental gradients. Future research should extend the network-MDE framework to other spatial gradients, including latitudinal patterns in network structure, to assess whether similar geometric constraints operate at larger scales. Additionally, incorporating species-specific environmental preferences, trait-based specialization, and habitat heterogeneity into null models will help refine predictions and improve our understanding of the processes that shape ecological networks along environmental gradients.

## Acknowledgements

We were supported by the Czech Science Foundation (19-14620S for PF and TMF, and 20-16499S and 21-24186M for RT, SS, ŠJ, INK, and AS). We are grateful to Robert Colwell for feedback on the manuscript.

