## Supplementary materials for "Does the mid-domain effect shape interaction networks along environmental gradients?"

**Table S1**. Summary of simulation model parameters, their potential values and ranges of simulation scenarios.

| **Model parameter** | **Value or range** |
|---|---|
| Artificial gradient (ENVI) | Bounded domain of 100 arbitrary units (1–100) |
| Number of lower-level species (LO) | 100 |
| Number of higher-level species (HI) | 200 |
| Lengths of species ranges (range length) | Three scenarios: uniform distribution of 1–100, 1–50, and 1–25 |
| Species range midpoints randomly placed along the gradient | Uniform 1–100 |
| higher-level species degree (number of interaction partners) | Three categories: specialists (2–20), intermediate specialists (20–50), and generalists (50–90) |
| Specialization along the gradient | Two scenarios: random, increasing |
| Gradient bin length | 10 bins of 10 arbitrary units each (1–10, 11–20, etc.) |
| Number of replicated simulations | 100 |

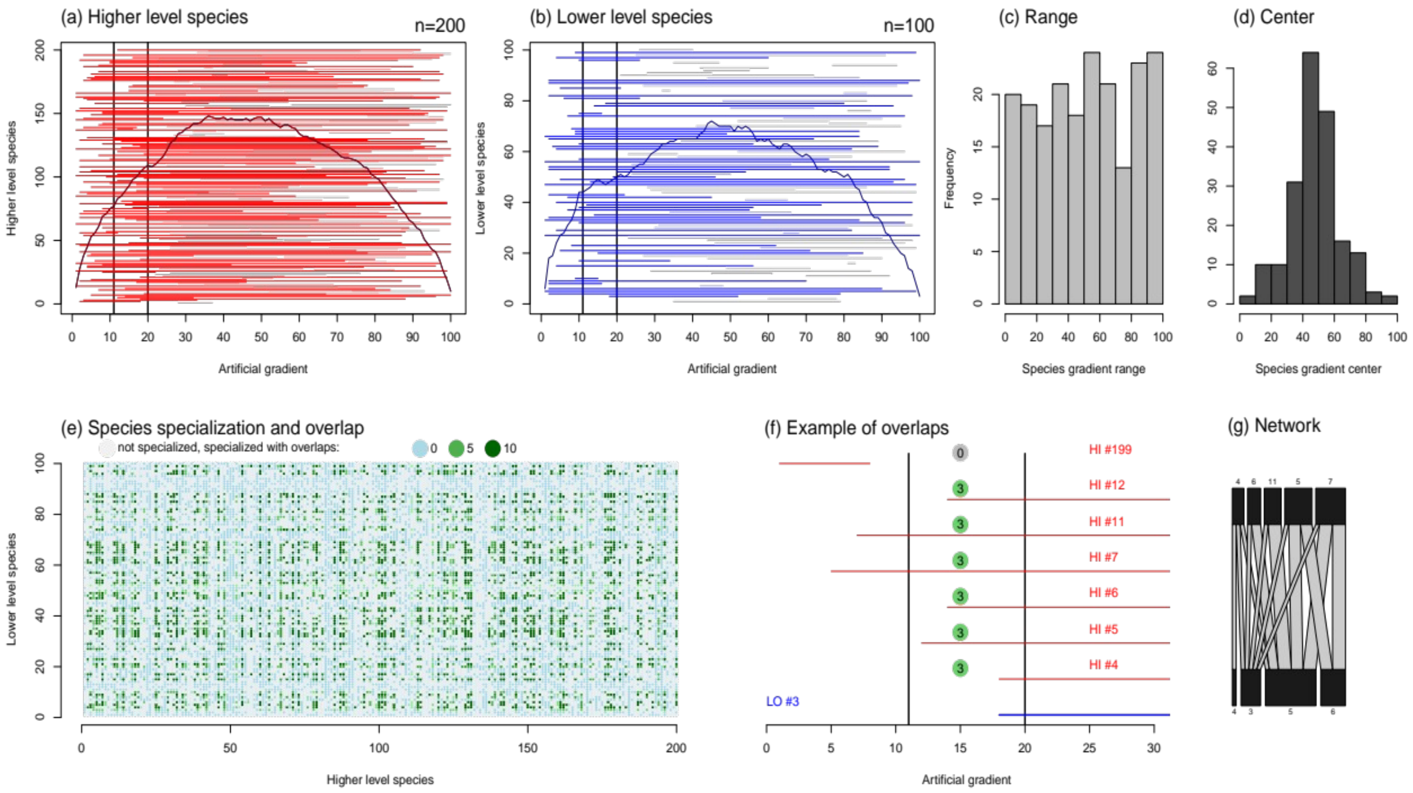


**Fig. S1 Illustration of the null modelling process for a given bin.** Distributions of simulated (a) higher- and (b) lower-level species ranges along the gradient, with the species ranges overlapping with the 11–20 bin highlighted in red or blue, species not in the bin are greyed out. The distributions of species ranges (c) and their centres (d) are shown by histograms. The resulting interaction network (e), specifically from the 11–20 bin. Each pair of species is coloured by whether it was modelled to potentially interact; grey indicates no potential interaction between the pair, light blue indicates that the pair can potentially interact, but the species do not overlap within this particular bin, and green indicates that the pair interacts within the bin (with darker greens denoting higher interaction frequency). Note the high proportion of non-interacting pairs; either because they were simulated as not potentially interacting (grey), or despite the pair potentially interacting, one species of the pair was not present in the 11–20 bin, or both species were present but their ranges within the bin did not overlap (light blue). (f) The calculation of interaction strengths between the lower-level species “LO #3” and seven upper level species in the 11–20 bin. The numbers in the circles are interaction strengths, calculated as the length of overlap between the upper-level species and the lower-level species within the gradient bin. (g) An illustrative bipartite network between higher and lower-level species.

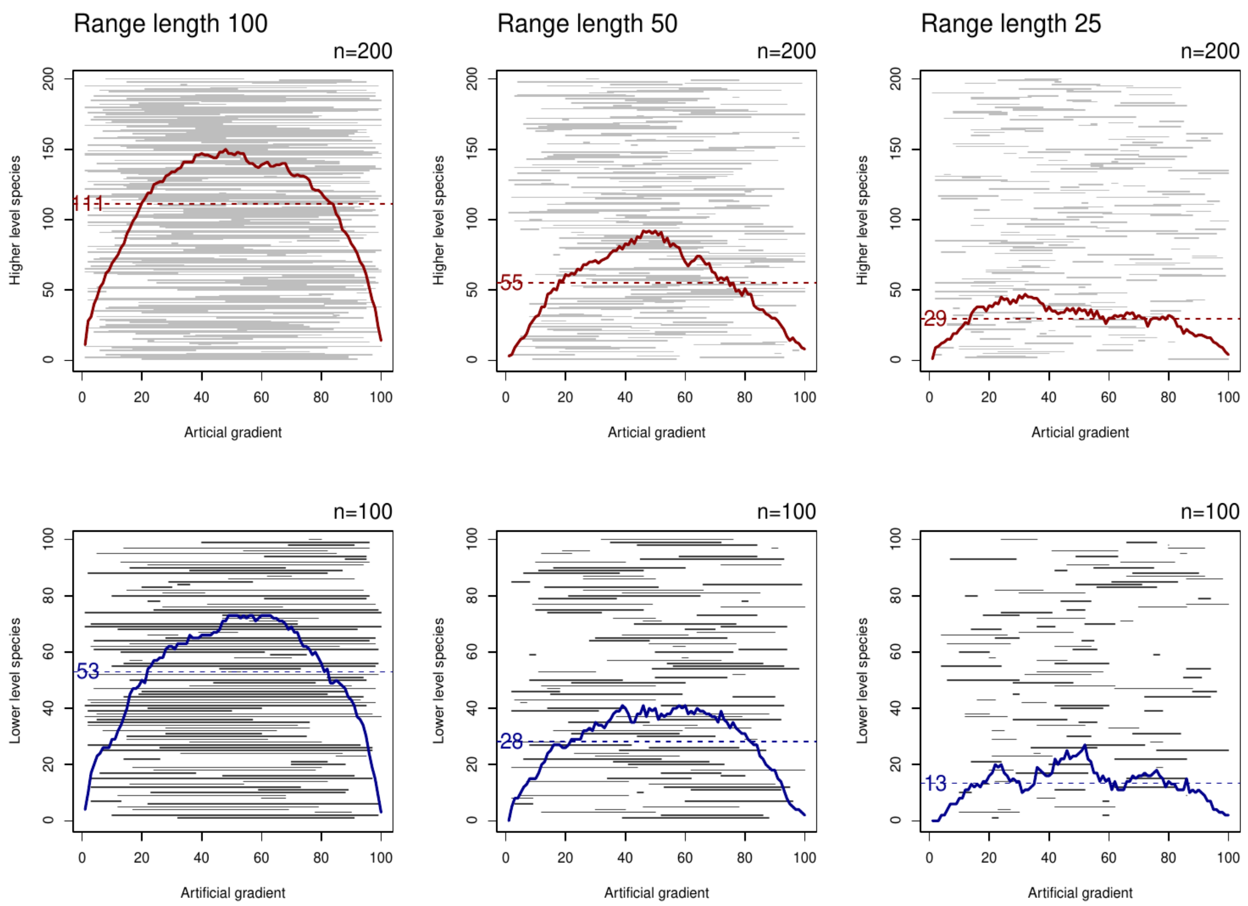


**Fig. S2** Patterns of higher- (top) and lower-level (bottom) species richness along the artificial gradient (1–100) for different lengths of species ranges (in columns). Gray lines correspond to species ranges. Coloured lines show overlaps of species ranges, therefore corresponding to species richness along the gradient (solid line). The mean number of species per combination is marked with a dashed line.

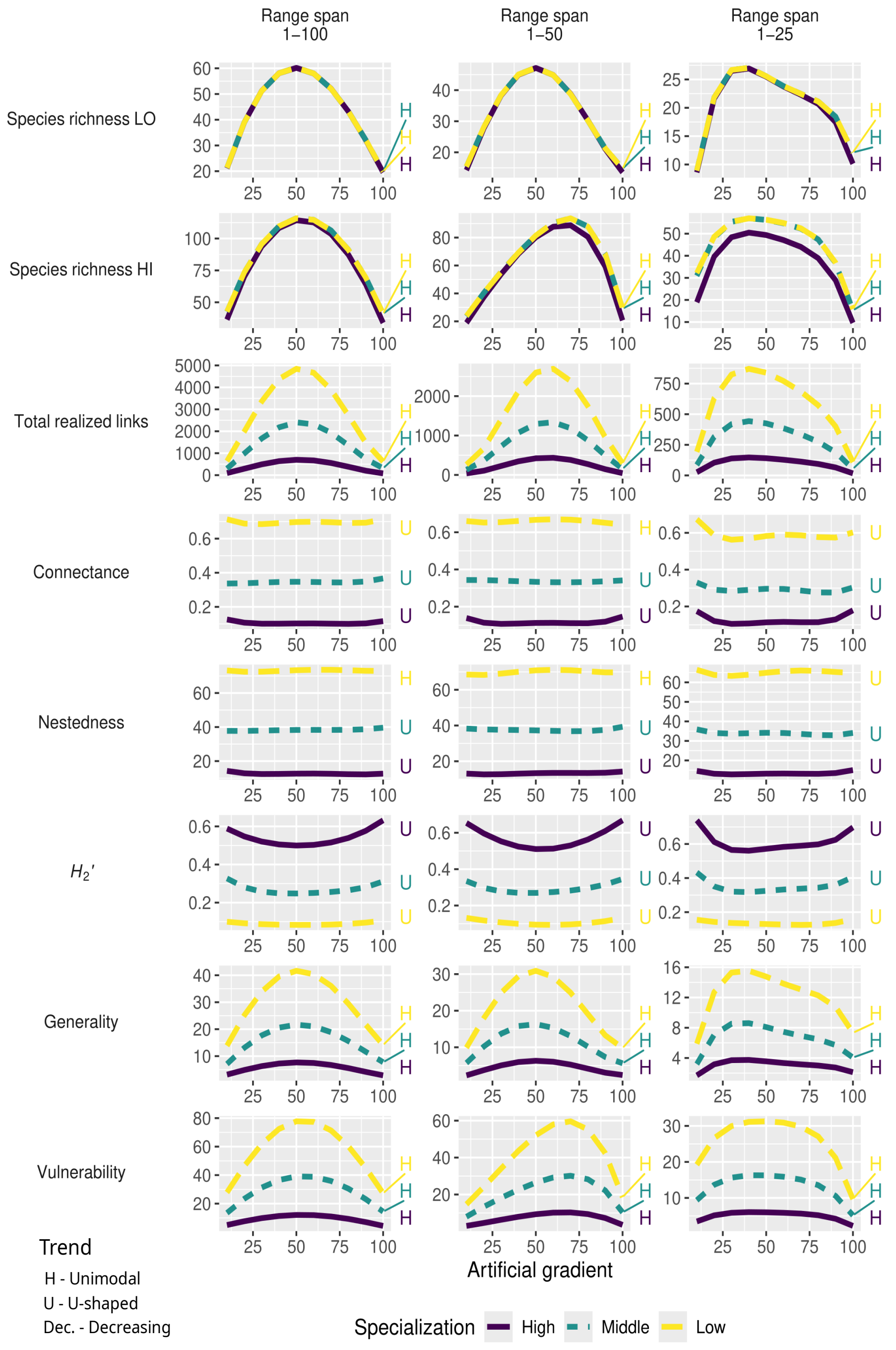


**Fig. S3** Predicted changes in network structure under mid-domain models where higher-level species-specialisation (degree) is *randomly* distributed along the artificial gradient. Panel columns represent modelled scenarios with different species' range sizes, from wide-ranged (left) to short-ranged (right) species. Panel rows correspond to network metrics. Within each panel, the best-fitting models are shown for simulations with the three categories of modelled species degree: yellow dashed lines represent low specialization with generalists in the community (species degree 50–90), green short-dashed lines represent middle specialization with intermediate specialists (20–50), and purple solid lines represent high specialization with specialists (2–20). The *fourth-degree* polynomial trends of individual lines are described by letters: H – unimodal (hump-) shaped, and U – U-shaped.

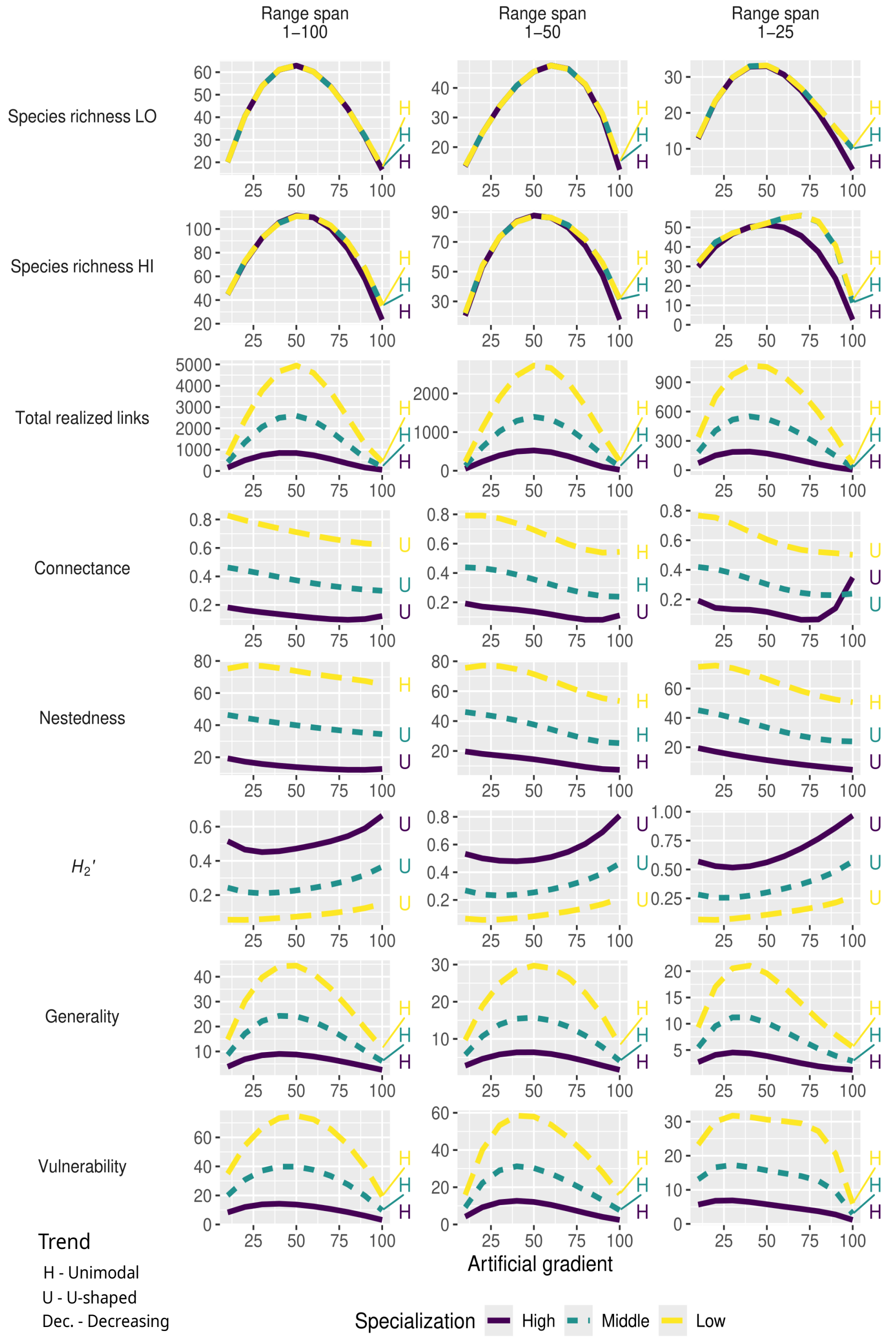


**Fig. S4** Predicted changes in network structure under mid-domain models where higher-level species-specialisation (degree) systematically increases along the artificial gradient. Panel columns represent modelled scenarios with different species' range sizes, from wide-ranged (left) to short-ranged (right) species. Panel rows correspond to network metrics. Within each panel, the best-fitting models are shown for simulations with the three categories of modelled species degree: yellow dashed lines represent low specialization with generalists in the community (species degree 50–90), green short-dashed lines represent middle specialization with intermediate specialists (20–50), and purple solid lines represent high specialization with specialists (2–20). The *fourth-degree* polynomial trends of individual lines are described by letters: H – unimodal (hump-) shaped, U – U-shaped, Dec. – linearly decreasing, and Inc. – linearly increasing.

**Text S1**. Further details about the four empirical networks.

The tropical myrmecophytic ant–plant dataset was collected in a wet primary rainforest on the slopes of Mount Wilhelm, Papua New Guinea (5° 43’ 18’’ S, 145° 16’ 12’’ E). Sampling was conducted at nine elevations spanning a gradient from 700 to 1600 m a.s.l, with 100 m intervals between sites. At each sampling location, all understorey trees (up to 15 m height) were checked for entrance holes and ant activity in stems, branches, or other pre-formed domatia, and all ant-inhabited trees were tagged. Ants were collected from trees with observed ant activity, and identified to morphospecies and species where possible, using existing reference collections and DNA barcoding (see Plowman *et al.* 2017 for more details).

The two tropical plant–pollinator datasets were collected along a continuous elevational gradient in primary rainforest on Mount Cameroon, Cameroon (4° 12′ 10″ N, 9° 10′ 12.0″ E), spanning four elevations from 650 to 2200 m a.s.l. Data were sampled during dry and wet seasons. The temperate plant-pollinator dataset was collected along an elevational gradient of semi-natural forests in the Krkonoše Mountains, Czechia (50° 36′ 48.33″ N, 15° 46′ 40.65″ E), at four elevations ranging from 450 to 1000 m a.s.l. The same sampling protocol was followed for all plant–pollinator datasets. At each sampled elevation, six transects (200 m × 10 m) were established to cover local diversity of flowering plants. Video cameras filmed specimens (five in Cameroon and seven in Krkonoše) of all flowering plant species for 24 hours to record flower visitors. The recorded footage was processed post-fieldwork to identify visitors to species or classify into morphospecies by experienced entomologists, and their behaviours were categorised to determine whether they acted as potential pollinators (further details in Klomberg *et al.* 2020; Sakhalkar *et al.* 2023). Only visitors identified into species or morphospecies reliably recognisable across elevations were included in our analyses. Species number in the empirical networks are show in **Fig. S5**.

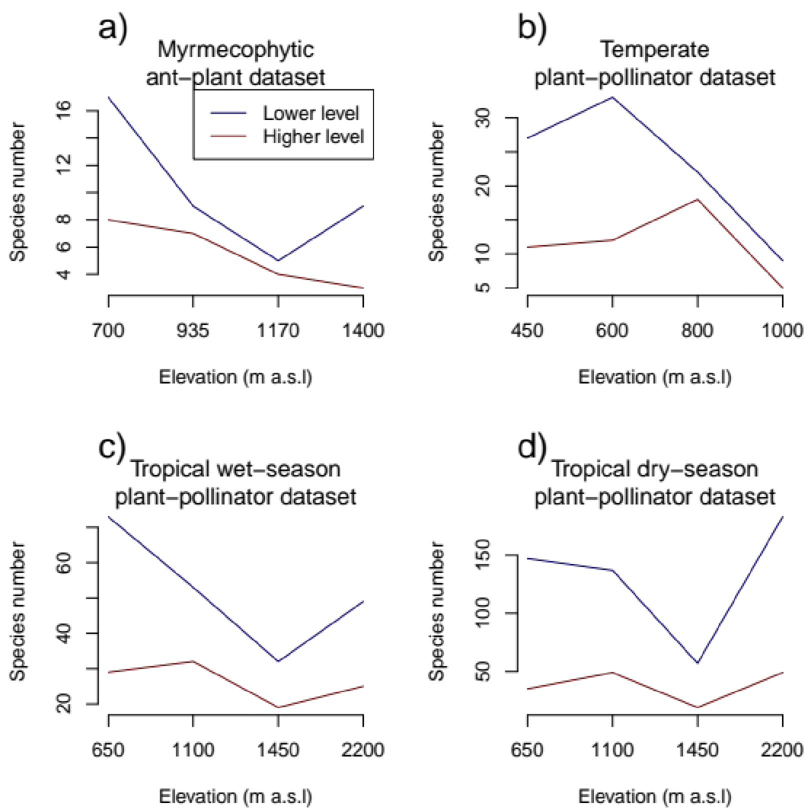


**Fig. S5** Higher- and lower-level number of species along the elevation for each of the empirical networks

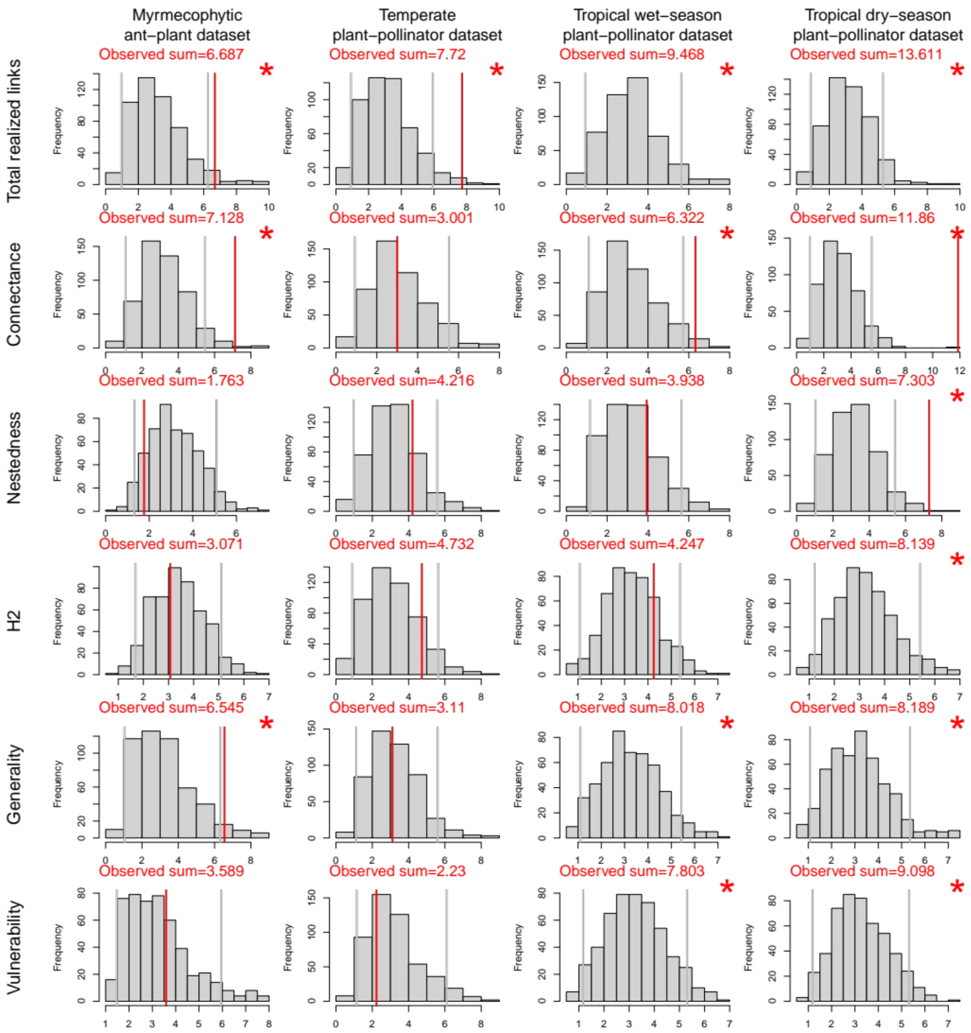


**Fig. S6** Histograms of sums of standardised effect sizes (SES) from observed (red line) and randomized (grey bars) interaction networks from **Fig. 5**. Horizontal grey lines define the 95% range of the sums of SES from randomized networks. * denotes instances where the sum of the observed SES is in the highest 5% of the distribution of the sums of SES from randomized networks.
